# Automated Optimization of Residual Reduction Algorithm Parameters in Opensim

**DOI:** 10.1101/2021.10.06.463431

**Authors:** Jordan T. Sturdy, Anne K. Silverman, Nathan T. Pickle

## Abstract

The residual reduction algorithm (RRA) in OpenSim is designed to improve dynamic consistency of kinematics and ground reaction forces in movement simulations of musculoskeletal models. RRA requires the user to select numerous tracking weights for the joint kinematics to reduce residual errors. Selection is often performed manually, which can be time-consuming and is unlikely to yield optimal tracking weights. A multi-heuristic optimization algorithm was used to expedite tracking weight decision making to reduce residual errors. This method produced more rigorous results than manual iterations and although the total computation time was not significantly reduced, this method does not require the user to monitor the algorithm’s progress to find a solution, thereby reducing manual tuning. Supporting documentation and code to implement this optimization is freely provided to assist the community with developing movement simulations.

## 1. INTRODUCTION

Musculoskeletal models and simulations have seen significant increase in adoption in recent years, facilitated by software platforms such as OpenSim (Delp et al., 2007, simtk.org). Inverse dynamics calculations are frequently the basis of these simulations. Inverse dynamics uses experimental observations, i.e., ground reaction forces and joint kinematics, to calculate the joint moments required to produce these observed data (Yamaguchi, 2005). This calculation contains multiple sources of error including modeling assumptions (e.g., rigid body assumptions, segment inertial properties), and measurement error (e.g., soft tissue artifact). These sources of error result in dynamic inconsistency (Kingma and Toussaint, 1996; Pearsall and Costigan, 1999; Riemer et al., 2008), meaning additional fictitious forces are required to produce a simulated motion that replicates the experimental data. Several methods have been proposed to reduce errors to improve dynamic consistency (Cahouët et al., 2002; Ganley and Powers, 2004; Kuo, 1998), including OpenSim’s residual reduction algorithm (RRA, https://simtk-confluence.stanford.edu/display/OpenSim/Residual+Reduction+Algorithm) that reduces the fictitious, or “residual”, forces and moments applied at the pelvis. RRA first computes changes to the total model mass as Δ*m = F*_*y,avg*_*/g*, where *F*_*y,avg*_ is the average vertical residual force across the simulation and *g* is acceleration due to gravity. RRA also calculates an altered torso center of mass (COM) location to reduce residual moments in the frontal and transverse planes. In addition, RRA can change the model joint accelerations slightly to achieve better dynamic consistency and further reduce residual forces and moments. Thus, there is a tradeoff between the residuals and the kinematic errors, where the tradeoff is governed by user-selected tracking weights. OpenSim’s documentation recommends users manually adjust the tracking weights until the residuals are minimized, which is a time-consuming process. In this paper we describe an open-source software tool (provided in Python and MATLAB) for optimization of tracking weights in RRA, which provides biomechanics researchers with an automated method for selecting RRA tracking weights, increasing efficiency in developing high-quality movement simulations.

## 2. METHODS

A Tracking Weight Selection Algorithm (TWSA) was developed to optimize the tracking weight values, which serve as inputs to the optimization algorithm RRA in OpenSim. A modified random hill climbing algorithm was used to explore the search space.

### 2.1. OBJECTIVE FUNCTION FORMULATION

The TWSA minimized the sum of root-mean-squared (RMS) residual forces and moments and the sum of RMS kinematic tracking errors as a weighted multi-objective cost function:

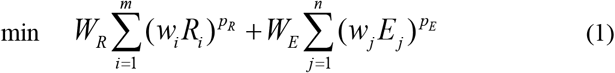

where *R*_*i*_ is the RMS of a residual term, *W*_*R*_ and *W*_*E*_ are the user-specified weights applied to the sums of the forces and errors, *w*_*j*_ and *w*_*i*_ are the weights applied to individual errors and forces, *E*_*j*_ is the RMS of a particular error, *p*_*R*_ and *p*_*E*_ are user specified powers and *m* and *n* are the number of residuals and errors examined respectively. Each RMS value is weighted by a user-specified value, *w*_*i*_ or *w*_*j*_. By default these weights are set to the inverse of 2cm for translational errors, 2° for rotational errors, 7.5% bodyweight (N) for residual forces, and 1.5% bodyweight times one meter for residual moments in line with OpenSim recommendations (Hicks et al., 2015 and OpenSim Documentation^1^). Thus, the weights serve to normalize the RMS errors to a desired threshold value. Errors above the threshold result in a weighted value >1, while errors below the threshold result in weighted values < 1. The exponents *p*_*R*_ and *p*_*E*_ are applied to the residuals and errors to cause errors above the threshold to rapidly increase the objective function value, while errors below the threshold diminish in the objective function.

### 2.2. ALGORITHM DESCRIPTION

The TWSA minimized the residual forces and moments while tracking the experimental data. A set of uniform tracking weights was used to perform the initial RRA and the objective function was evaluated in a MATLAB function and reported to the TWSA. This first evaluation of the objective function was the current best solution. An array of parameters, 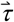, was then created to perturb tracking weights. Each element in 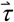 was calculated as *τ*_*i*_ = *b*^*t*^, where *b* started at 1.5 and was lowered to 1.1 after 75% of the TWSA iterations were performed, and *t* was an integer ranging from -2 to 2 selected from a random sample which was weighted to favor negative values (decrease the tracking weight) if coordinate tracking was below a good threshold, positive values if tracking was bad (increase the tracking weight), and equal chances otherwise. Each tracking weight was then multiplied by the corresponding *τ*_*i*_ to generate a new set of tracking tasks, which were used to run the next RRA iteration. The objective function was evaluated after each iteration, and the tracking weights corresponding to the current lowest objective function value were used as the starting weights that are perturbed for subsequent iterations. This process was repeated until the cost function value was below a specified threshold or an iteration limit set by the user was reached. The resulting solutions from the TWSA do not guarantee optimality but improve errors relative to baseline, meaning that the solutions have lower residual forces and moments while still maintaining acceptable kinematic tracking errors (Hicks et al., 2015).

### 2.3. TEST CASE

To test the effectiveness of the TWSA, we selected a single participant (subject01 - male, 72.84kg) from publicly available data repository (simtk.org/home/nmbl_running, Hamner & Delp, 2013) for a common dynamic task: running. Three running strides each were isolated from motion files of 2.0, 3.0, 4.0, and 5.0 m/s, and were used to develop the running simulations in OpenSim 3.3. The TWSA was applied using two different models. First, the full body model with 29 degrees of freedom and 92 Hill-type musculotendon actuators developed by Hamner et al. (Hamner et al., 2010; Hamner and Delp, 2013) was used because it was consistent with the dataset. Next, the gait2392 model with 23 degrees of freedom and 92 Hill-type musculotendon actuators (Delp et al., 2007) was selected because it is widely used and available with OpenSim installation. A key distinction between these models is that gait2392 uses a lumped head/arms/trunk segment, whereas the Hamner2010 model implements multi-segment arms. This difference in model formulation can be used to demonstrate the robustness of the TWSA, as ignoring contributions of the arms during running introduces additional dynamic inconsistency in the simulation. Following a typical workflow with ground reaction forces applied to the calcanei and tracking a desired inverse kinematics solution, RRA was first run iteratively using fixed tracking weights, and recommended model mass adjustments were performed to reduce average residual forces and moments applied to the pelvis until the total mass change was less than 0.001 kg. Then, the TWSA was applied to each running trial using the mass adjusted model to further reduce the magnitude of residuals. For this study, a fixed iteration number of 200 RRA iterations were performed by the TWSA, and the tracking weights associated with the best objective function value were then re-run for the final RRA results. For all trials *W*_*E*_ = 3, *W*_*R*_ =1, *p*_*R*_ = 3, and *p*_*E*_ = 4. Residuals after the TWSA were compared to residuals prior to the TWSA, but after the mass adjustments, to determine simulation improvement. The peak net external force from each running trial was used to normalize residuals for evaluation (Hicks et al., 2015), and both peak and RMS residuals were analyzed. Tracking errors were evaluated to ensure all coordinates were within measurement error.

## 3. RESULTS

Mass changes were less than 0.5 kg for all trials using both the Hamner2010 model and the gait2392 model. Torso center of mass changes were less than 6 cm for most trials, and adjustments were slightly larger for the Hamner2010 model (Table 1). Trial1 in the 2.0 m/s condition required the largest change in center of mass location (>10 cm for both models) during this portion of the conventional RRA workflow.

**Table 1.** Changes made to the torso center of mass for each trial using both the Hamner2010 and gait2392 models. Values are reported in cm as (x, z) pairs for the anterior (x) and medio-lateral (z) directions.

|  |  | Hamner2010 |  |  | gait2392 |  |  |
| --- | --- | --- | --- | --- | --- | --- | --- |
|  |  | Trial |  |  | Trial |  |  |
|  |  | 1 | 2 | 3 | 1 | 2 | 3 |
| Speed<br>(m/s) | 2.0 | (15.7, -1.2) | (4.4, 1.3) | (5.9, 0.3) | (11.8, -0.7) | (2.8, 1.2) | (4.2, 0.4) |
|  | 3.0 | (5.0, 1.3) | (3.6, 0.9) | (4.8, 1.0) | (3.5, 1.4) | (2.3, 1.0) | (3.1, 1.2) |
|  | 4.0 | (3.5, 1.4) | (5.8, 1.6) | (4.6, 0.9) | (2.1, 1.5) | (4.0, 1.5) | (3.1, 1.0) |
|  | 5.0 | (9.0, 2.2) | (4.9, 0.5) | (7.9, -0.4) | (6.3, 1.9) | (3.3, 1.1) | (5.6, -0.4) |

After performing the standard RRA-based model adjustments, we ran the TWSA using MATLAB R2020a (The Mathworks, Inc.) and OpenSim v3.3 (Delp et al., 2007, simtk.org) on a 3.60 GHz CPU with 12 logical processors to optimize tracking weights, with each running speed condition (2.0, 3.0, 4.0, and 5.0 m/s) evaluated in parallel. The 200 iterations took the most time to complete during the 2 m/s condition, finishing in 2 hours and 4 minutes using the Hamner2010 model, and 49 minutes using the gait2392 model. TWSA objective function values were substantially reduced in the first 50 RRA iterations and achieved reasonable convergence between 100 and 150 iterations (Fig. 1). However, gradual reductions continue to occur with more iterations, and the lowest objective function values were produced in the last 50 iterations for all trials. Using both the Hamner2010 and gait2392 models, the TWSA produced objective function values less than 2.5 on average for 3, 4 m/s trials. Average best objective function values were 8.99 and 5.84 at 2 and 5 m/s, respectively, using the Hamner2010 model and were 5.28 and 8.26 using the gait2392 model at 2 and 5 m/s, respectively. Trial 1 at 2 m/s had the highest objective function value for both models, but the 2 m/s trials otherwise achieved values less than 1.1. In addition, certain trials sporadically produced peaks in objective function values between 50 and 150 iterations of the TWSA, and these were most pronounced using the Hamner2010 model (Fig. 1).

**Figure 1.**
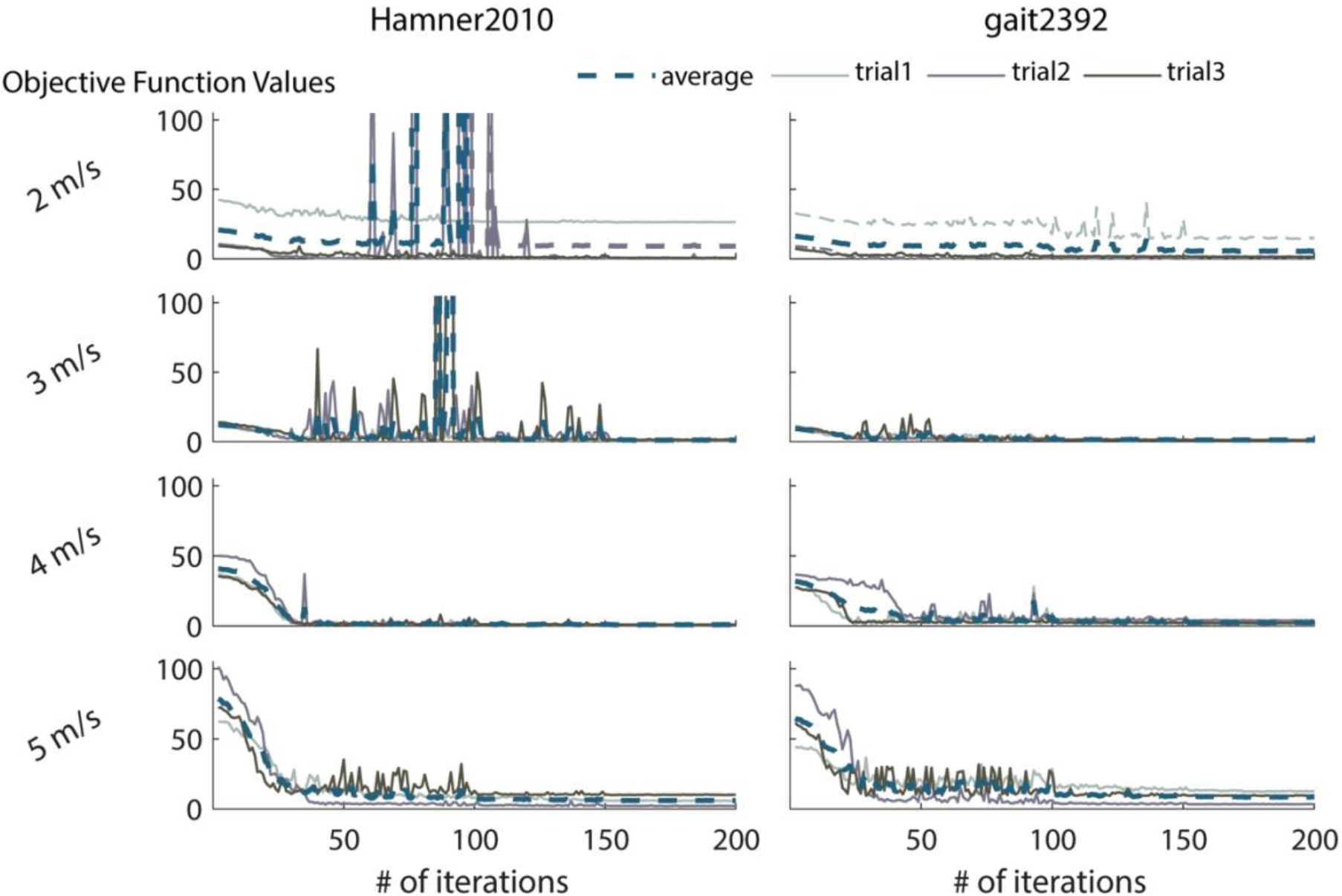
Objective function values for the tracking weight selection algorithm optimization at each RRA iteration using the Hamner2010 model (left) and gait2392 model (right). Three trials were averaged at each iteration for both models at all running speeds (blue dashed lines).

After tracking weight optimization, peak and RMS residuals were substantially reduced except for the FZ residual (Fig. 2). RMS forces were less than 1% peak external force and RMS moments were less than 0.35% peak external force times 1 meter for both models after tracking weight optimization when averaged across speeds, with improvement in all residuals except FZ (Table 2.). Residuals were slightly larger using the gait2392 model compared to the Hamner2010 model after tracking weight selection. In addition, the FY residual was improved more across all speeds when using the Hamner2010 model compared to gait2392 achieving both a lower peak (Fig 2.) and RMS value (Table 2).

**Table 2.** Root mean squared (RMS) residual forces and moments normalized to OpenSim recommended maximum values (Hicks et al., 2015). RMS forces and moments are normalized to 5% and 1% respectively of the peak external force (net ground reaction force) during each trial. Values reported are the average RMS of 3 trials for each running speed both before (Pre TWSA) and after (Post TWSA) the tracking weight optimization using both models. Values above 1 indicate the residual exceeds recommended thresholds.

|  |  | Hamner2010 |  |  |  |  |  | gait2392 |  |  |  |  |  |
| --- | --- | --- | --- | --- | --- | --- | --- | --- | --- | --- | --- | --- | --- |
|  |  | FX | FY | FZ | MX | MY | MZ | FX | FY | FZ | MX | MY | MZ |
| Pre<br>TWSA | 2.0 m/s | 0.44 | 0.27 | 0.13 | 2.66 | 3.84 | 3.45 | 0.40 | 0.24 | 0.11 | 3.30 | 2.98 | 3.29 |
|  | 3.0 m/s | 0.31 | 0.18 | 0.11 | 2.45 | 4.67 | 2.48 | 0.26 | 0.16 | 0.08 | 2.89 | 3.61 | 2.55 |
|  | 4.0 m/s | 0.41 | 0.20 | 0.14 | 3.03 | 4.78 | 2.42 | 0.35 | 0.19 | 0.08 | 3.19 | 2.91 | 2.30 |
|  | 5.0 m/s | 0.52 | 0.25 | 0.18 | 3.30 | 6.00 | 3.01 | 0.46 | 0.23 | 0.13 | 3.58 | 2.94 | 3.01 |
| Post<br>TWSA | 2.0 m/s | 0.24 | 0.11 | 0.11 | 0.80 | 0.11 | 0.17 | 0.22 | 0.18 | 0.09 | 0.64 | 0.87 | 0.06 |
|  | 3.0 m/s | 0.12 | 0.11 | 0.12 | 0.02 | 0.29 | 0.15 | 0.13 | 0.14 | 0.08 | 0.07 | 0.17 | 0.37 |
|  | 4.0 m/s | 0.04 | 0.07 | 0.15 | 0.21 | 0.24 | 0.16 | 0.15 | 0.16 | 0.08 | 0.27 | 0.26 | 0.24 |
|  | 5.0 m/s | 0.23 | 0.13 | 0.17 | 0.19 | 0.32 | 0.19 | 0.26 | 0.24 | 0.12 | 0.17 | 0.08 | 0.42 |

**Figure 2.**
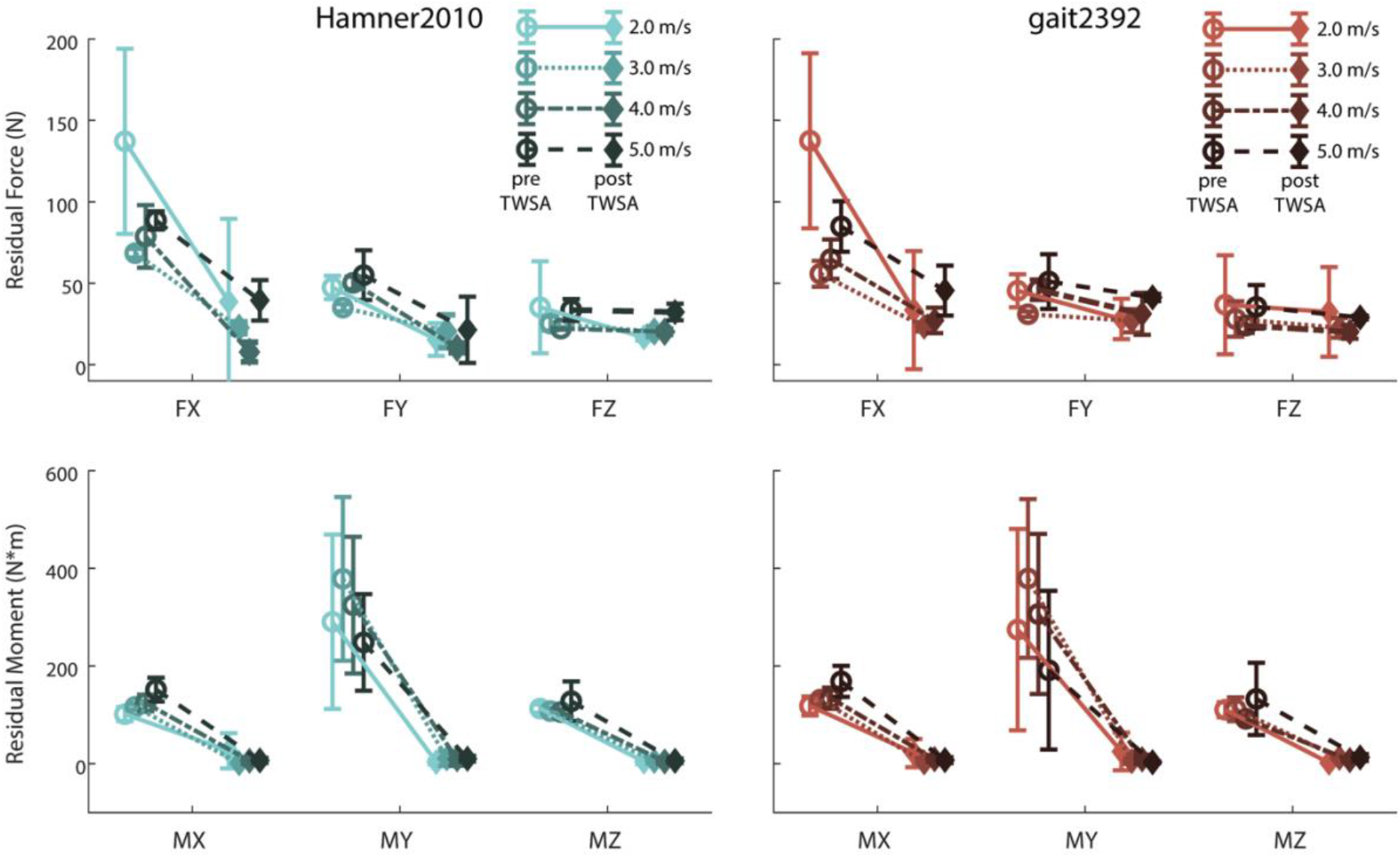
Peak residual forces (top) and moments (bottom) averaged at each running speed and plotted before (pre) and after (post) the tracking weight selection algorithm was applied. Error bars represent the standard deviation of three trials at each speed using the Hamner2010 (left) and gait2392 (right) models.

## 4. DISCUSSION

The TWSA performed well using two different musculoskeletal models and improved residual forces and moments by similar amounts with each model. We performed 200 iterations of RRA within the TWSA, but this upper limit may not be necessary to obtain comparable improvements in residuals. This upper limit can also be adjusted by the user. Other than for one trial in both 2 and 5 m/s using the gait2392 model, the best objective function value was improved by less than 1 in the last 100 RRA iterations. While more iterations will guarantee a broader search of the solution space, a “good enough” objective function value could be used as a threshold to stop the TWSA regardless of the number of iterations performed. An objective value threshold implementation, also an option within the tool, would allow researchers to use more computational time on trials with higher residuals, like the first 2 m/s trial in this study.

All residual moments were substantially improved by the TWSA, while improvements in residual forces differed between models and directions. Some of this discrepancy between residuals may be attributed to the objective function formulation. The recommended normalization for residuals is stricter for moments than for forces, and thus, the TWSA favors solutions with lower moment residuals. In addition, the RMS residual forces were all below the recommended threshold used to weight the objective function contributions and were not highly penalized. Force residuals in the Z direction were not substantially reduced for either model at any running speeds. Z (medial/lateral) direction forces were lower than both X (anterior/posterior) and Y (vertical) forces prior to the TWSA, and thus contributed less to objective function values compared to X and Y forces. Improvements in Y direction force residuals were greater using the Hamner2010 model compared to gait2392. This difference suggests that the presence of arm swing dynamics in the Hamner2010 model was leveraged by the TWSA to reduce residual vertical forces without introducing excessive kinematic tracking errors. The residual forces and moments are trivial to reduce if the kinematics are not tracked, which results in very low residuals but very high errors. The TWSA sought to balance reducing the residuals and reducing kinematic error. Trial1 at 2 m/s illustrates how this balance of terms may limit the ability of the TWSA to achieve a low cost function value. Kinematic errors likely required greater residuals reductions during this trial, and these errors are much larger than the weights used for normalization. The user must choose whether to allow greater kinematic errors or accept the large residuals in this case and adjust weights accordingly, and output results should be critically examined before implementing more in-depth simulation analyses.

Other algorithms have been used to automate RRA tracking weight selection using similar objective functions (e.g., Samaan et al., 2016), and many different optimization approaches could be successfully applied to this application. While it is challenging to directly compare the TWSA to others because different motions were used and cost function formulations are different, the kinematic accuracy of our approach (see Appendix for full kinematic results) was comparable to other approaches (Samaan et al., 2016).

## 5. CONCLUSIONS

The TWSA provides a valuable new tool for efficiently computing optimal RRA tracking weights. This algorithm is an important and practical contribution to the modeling and simulation community and may be used to improve the quality of musculoskeletal simulations of human movement. The TWSA reduced residual forces and moments below established guidelines in a reasonably short amount of time (<2hr), requiring little or no user input. Supporting documentation and code to implement the TWSA in both Python and MATLAB are available for download at: https://github.com/FxnlBiomechLab/twsa-optimization-for-rra.

## ACKNOWLEDGEMENTS

The authors would like to thank Daniel Anderson and Moriah Hunt for their contributions to the development of the initial TWSA algorithm. The authors also acknowledge Amy Hegarty and Alexandra Newman for testing of the tool.

## APPENDIX

**Table A1.** Largest root mean squared tracking error for each coordinate across all running speeds and trials using the Hamner2010 model.

| Hamner2010 |  |  |  |  |  |  |
| --- | --- | --- | --- | --- | --- | --- |
| Translational<br>(cm) |  | pelvis_tx | pelvis_ty | pelvis_tz |  |  |
|  | Pelvis | 0.92 | 1.14 | 0.97 |  |  |
| Rotational<br>(deg) | Pelvis | pelvis_tilt | pelvis_list | pelvis_rotation |  |  |
|  |  | 0.54 | 1.24 | 1.96 |  |  |
|  | Torso | lumbar_extension | lumbar_bending | lumbar_rotation |  |  |
|  |  | 1.61 | 2.28 | 2.16 |  |  |
|  | Right leg | hip_flexion | hip_adduction | hip_rotation | knee_angle | ankle_angle |
|  |  | 1.99 | 1.19 | 0.19 | 0.85 | 0.19 |
|  | Left leg | hip_flexion | hip_adduction | hip_rotation | knee_angle | ankle_angle |
|  |  | 1.10 | 1.08 | 0.76 | 0.30 | 0.17 |
|  | Right arm | arm_flex | arm_add | arm_rot | elbow_flex | pro_sup |
|  |  | 0.38 | 0.80 | 0.13 | 0.53 | 0.15 |
|  | Left arm | arm_flex | arm_add | arm_rot | elbow_flex | pro_sup |
|  |  | 1.07 | 0.65 | 0.73 | 0.23 | 0.01 |

**Table A2.** Largest root mean squared tracking error for each coordinate across all running speeds and trials using the gait2392 model.

| Gait2392 |  |  |  |  |  |  |
| --- | --- | --- | --- | --- | --- | --- |
| Translational<br>(cm) |  | pelvis_tx | pelvis_ty | pelvis_tz |  |  |
|  | Pelvis | 1.12 | 0.69 | 1.05 |  |  |
| Rotational<br>(deg) | Pelvis | pelvis_tilt | pelvis_list | pelvis_rotation |  |  |
|  |  | 0.71 | 0.35 | 0.44 |  |  |
|  | Torso | lumbar_extension | lumbar_bending | lumbar_rotation |  |  |
|  |  | 1.66 | 1.00 | 0.85 |  |  |
|  | Right leg | hip_flexion | hip_adduction | hip_rotation | knee_angle | ankle_angle |
|  |  | 0.48 | 0.45 | 0.29 | 0.61 | 0.06 |
|  | Left leg | hip_flexion | hip_adduction | hip_rotation | knee_angle | ankle_angle |
|  |  | 1.92 | 0.72 | 0.17 | 0.09 | 0.04 |

## Footnotes

1 https://simtk-confluence.stanford.edu:8443/display/OpenSim/Getting+Started+with+RRA

